# Deletion of genes encoding PU.1 and Spi-B leads to B cell acute lymphoblastic leukemia associated with driver mutations in Janus Kinases

**DOI:** 10.1101/291054

**Authors:** Carolina R. Batista, Michelle Lim, Anne-Sophie Laramée, Faisal Abu-Sardanah, Li S. Xu, Rajon Hossain, Rodney P. DeKoter

## Abstract

Precursor B-cell acute lymphoblastic leukemia (B-ALL) is associated with recurrent mutations that occur in cancer-initiating cells. There is a need to understand how spontaneous driver mutations influence clonal evolution in leukemia. The ETS-transcription factors PU.1 and Spi-B (encoded by *Spi1* and *Spib*) execute a critical role in B cell development and serve as complementary tumour suppressors by opposing the proliferative events mediated by IL-7R signaling. Here, we used a mouse model to conditionally delete *Spi1* and *Spib* genes in developing B cells. These mice developed B-ALL with a median time to euthanasia of 18 weeks. We performed RNA and whole-exome sequencing (WES) on leukemias isolated from Mb1-CreΔPB mice and identified single-nucleotide variants (SNVs) in *Jak1*, *Jak3* and *Ikzf3* genes, resulting in amino acid changes and in the gain of early stop-codons. JAK3 mutations resulted in amino acid substitutions located in the pseudo-kinase (R653H, V670A) and in the kinase (T844M) domains. Introduction of these mutations into wild-type pro-B cells conferred survival and proliferation advantages. We conclude that mutations in Janus kinases represent secondary drivers of leukemogenesis in the absence of Spi-B and PU.1 transcription factors. This mouse model represents an useful tool to study clonal evolution and tumour heterogeneity in B-ALL.

## Introduction

Acute lymphoblastic leukemia is the most common type of childhood cancer, with approximately 6000 new cases diagnosed in the United States each year^1^. Most leukemias originate within the B cell rather than the T cell lineage^2,3^. Precursor B cell acute lymphoblastic leukemia (pre-B-ALL) is a disease that is revealed by the presence of transformed precursor B cells in the blood, bone marrow, and tissues; and is most common in 1-5 year old patients^4^. Most pre-B-ALL cases are associated with genetic abnormalities that include chromosomal translocations or point mutations. In pre-B-ALL, up to two thirds of genes with point mutations encode transcriptional regulators such as Pax-5, Ikaros, or EBF1^3^. Pre-B-ALL cells are frequently arrested at an early stage of development, express interleukin-7 receptor (IL7R), and have high levels of Janus Kinase (JAK)-STAT signaling to sustain survival and proliferation^5–7^. Activating mutations of the *IL7R* gene have been described in human pre-B-ALL^8^. *JAK* and *IL7R* mutations are frequent in several subtypes of pre-B-ALL including the recently described disease Ph-like leukemia^9,10^. In summary, mutations that activate cytokine signaling, and impair differentiation, function as driver mutations in pre-B-ALL.

PU.1 (encoded by *SPI1*) and Spi-B (encoded *SPIB* in mice) are transcription factors of the E26-transformation-specific (ETS) family^11^. These two proteins share a conserved DNA binding domain and interact with an overlapping set of DNA binding sites within the genome^12^. PU.1 and Spi-B complement one anothers function, and activate multiple genes involved in B cell receptor signaling^12–15^. Lack of these factors in developing B cells results in a block to B development at the small pre-B cell stage associated with impaired *Ig* light chain rearrangement^15,16^. Importantly, conditional deletion of Spi-B and PU.1 in developing B-cells leads to high incidence of B-ALL in mice, but the mechanisms of leukemogenesis in the absence of these transcription factors are still undetermined^17^. In conclusion, PU.1 and Spi-B are required for B-cell development, and function as complementary tumour suppressors in the B cell lineage.

B cell neoplasms, like all cancers, are thought to be diseases in which there is clonal evolution from a common precursor, in which acquired gene mutations drive an evolutionary natural selection process^18,19^. The mechanisms by which cancer-initiating cells respond to selection pressures during clonal evolution have been classified into a number of common hallmarks^20^. In response to selection pressure, the genetic makeup of cancer-initiating cells changes during the course of disease due to acquired mutation. Mutations can be broadly classified as drivers or passengers^18,21^. Driver mutations are those that provide a growth advantage to a cancer clone, whereas passenger mutations do not provide a growth advantage. Pediatric B-ALL is less curable upon relapse due to clonal evolution of the leukemia, resulting in driver mutations inducing a more aggressive disease^22^. High levels of intratumoral heterogeneity of mutations is a poor prognostic marker for leukemia^23^. Whole-exome or whole-genome sequencing of pre-B-ALL cases is expected to lead to a deeper understanding of the genetic causes of this disease, ultimately permitting molecular targeted therapy for individual patients^2^.

In this study, we investigated the molecular features of leukemogenesis in a spontaneous model of B-ALL induced by deletion of genes encoding PU.1 and Spi-B. We generated *Mb1*^+/Cre^*Spi1*^lox/lox^*Spib*^-/-^ mice, called here Mb1-CreΔPB mice^16^. We found that Mb1-CreΔPB mice developed pre-B-ALL characterized by the high expression of IL-7R, with a median time to euthanasia of 18 weeks. Using whole-exome sequencing (WES) and RNA-seq, we identified single-nucleotide variants (SNVs), most of which were predicted to have a role in the control of cell proliferation, communication and metabolism. Strikingly, we identified recurrent SNVs in genes encoding Aiolos, Jak1, and Jak3 in mouse leukemias. Further analysis revealed that SNVs located in *Jak3* resulted in three different types of amino acid substitutions within the pseudo-kinase domain (R653H, V670A) and kinase domain (T844M). We confirmed the ability of these mutations to provide survival and proliferation advantages to normal pro-B cells. In summary, this study shows that *Jak3* mutations are secondary drivers of leukemogenesis in the absence of Spi-B and PU.1. This mouse model may be useful to determine the effects of molecular targeted therapies on intratumoral heterogeneity and clonal evolution in B-ALL.

## Materials and Methods

### Mice and breeding

Mb1-Cre mice have been described previously^24^. For this study, Mb1-Cre mice were crossed with *Spi1*^lox/lox^ *Spib*^-/-^ to generate *Mb1*^+/Cre^ *Spi1*^lox/lox^ *Spib*^-/-^ mice (also referred to as Mb1-CreΔPB mice). C57BL/6 mice were purchased from Charles River Laboratories (Saint-Constant, QC, Canada). *Mb1^+^*^/Cre^*Spi1*^lox/lox^ *Spib*^+/+^ (referred to as Mb1-CreΔP) and *Mb1^+^*^/Cre^*Spi1*^+/+^ *Spib*^-/-^ or *Mb1^+^*^/+^Spi1^+/+^ *Spib*^-/-^ (referred to as Mb1-CreΔB) mice were used as experimental controls. Mice were fed with regular chow and tap water ad libitum and housed with a 12-h light–dark cycle. Mice showing signs of illness were euthanized and examined for spleen, thymus and lymph node enlargement and analyzed as described below. All experiments were performed in compliance with the Western University Council on Animal Care.

### Histology and microscopic analysis

Mb1-CreΔPB mice showing signs of disease were euthanized by CO_2_ and spleen and thymus were removed for histological analysis. Spleen and thymus removed from Mb1-CreΔB mice were used as control. Organs were fixed in 10% buffered formalin. Tissues were paraffin embedded, sectioned, and stained with hematoxylin and eosin. High-resolution micrographs were captured using a Q-Color3 digital camera (Olympus, Markham, ON, Canada).

### Flow Cytometry

Single cell suspensions were prepared from enlarged spleen and thymus from Mb1-CreΔPB mice. Red blood cells were removed from single-cell suspensions using hypotonic lysis. Flow cytometric analyses were performed using a FACSCanto or LSRII instrument (BD Immunocytometry Systems, San Jose, CA). Antibodies were purchased from eBioscience (San Diego, CA), BioLegend (San Diego, CA), or BD Biosciences (Mississauga, ON, Canada) and included PE-anti-CD19 (1D3), FITC-anti-BP-1 (6C3), APC-anti-IgM (II/ 41), PE-anti-Igk (187.1), FITC-anti-IL-7Rα (A7R34), BV421–anti-B220 (RA3-6B2), PE–anti-BP-1 (6C3). Data were analyzed using FlowJo 9.1 software (Tree Star, Ashland, OR).

### Whole-Exome Sequencing

Three matched-tumour and tail mouse samples (853, 854 and 857) had genomic DNA isolated using Wizard Genomic DNA Purification Kit (Promega Corporation, Madison WI). Whole-Exome Sequencing was performed by McGill University and Génome Québec Innovation Centre (Montreal, Canada). The SureSelectXT Mouse All Exon kit (Agilent Technologies, Mississauga, Canada) was used to perform exome target capture. Paired-end DNA library preparation was performed using SureSelectXT Target Enrichment System. An Illumina HiSeq4000 instrument was used to perform the whole-exome sequencing. BAM files were converted to Fastaq format on Galaxy suite^25^ using “SAMtoFastaq” tool. Fastaq reads had Illumina Adaptors were removed using TrimGalore!. Trimmed reads were aligned to the mouse reference genome (mm10) using Bowtie. BAM aligned files were used as input for single somatic variant callers. Somatic variants of three matched-tumour mouse samples were called by using two independent methods, Strelka and Mutect^26,27^. For both methods, variants were called using the standard settings recommend by the package documentation. VCF files containing the somatic nucleotide variants (SNVs) and small insertions and deletions (INDELs) generated by Strelka and Mutect were further filtered following the criteria ‘passed’ or ‘keep’ for the Strelka and Mutect outputs, respectively. Filtered VCF files outputted by Strelka and Mutect were intersected using VCFtools and annotated using the SnpEff^28^ package using the mouse genomic annotation (mm10). Annotated variants were classified according the variant impact in HIGH, MODERATE, MODIFIER and LOW, and overlapping variants among the three samples were determined. Independently, outputs generated by Strelka and Mutect were also annotated using the SnpEff tool for further analysis.

### RNA Sequencing

RNA was extracted from thymic tumours 853, 854 and 857 using RNeasy Kit (QIAGEN, ON, Canada). RNA-sequencing was performed by McGill University and Génome Québec Innovation Centre (Montreal, Canada). Paired-end (mRNA-sequencing stranded) libraries were prepared using Truseq stranded total RNA library prep kit. Libraries were sequenced using an Illumina HiSeq4000 PE100 instrument. Data analysis was performed as described previously^16^. In summary, Standard Illumina sequencing adaptors were removed using Trim Galore!. Trimmed Fastaq reads were aligned to the mouse reference genome version mm10 using TopHat2^29^ in mate-paired mode using Galaxy suite. Assembled transcript abundance was determined as previously using Cufflinks^29^.

### Sanger Sequencing

RNA extracted from 20 thymic leukemias was reversed transcribed to complementary DNA using iScript cDNA synthesis kit (Bio-Rad, Hercules, CA). Primers flanking the *Ikzf3*, *Jak3* and *Jak1* mutations identified by WES were designed using Primer-BLAST. cDNA was used as a template for PCR reactions using specific primers (Sequences described in **Supplementary Information**). PCR reactions were run in a 1.5% agarose gel and the PCR products were excised and purified from the gel using QIAquick Gel Extraction Kit (QIAGEN, ON, Canada). Purified DNA was submitted for Sanger sequencing at London Regional Genomics Centre, Western University (London, ON, Canada). Sequencing chromatograms and DNA sequences were analyzed in the software 4Peaks (Mekentosj, Amsterdam).

### DNA Constructs and Site-Directed Mutagenesis

The mouse MSCV-IRES-GFP-JAK3 plasmid was kindly provided by Dr. Kevin D. Bunting. T844M, R653H, and V670A mutations were generated by site-directed mutagenesis using Q5^®^ Site-Directed Mutagenesis Kit (New England BioLabs, Ipswich, MA). Site-directed mutagenesis primers for each one of the specific mutations were designed using NEBaseChanger tool. Mutations were confirmed by Sanger Sequencing before retroviral production and spinoculation.

### Retroviral production and infection

Retroviral supernatants for wild-type Jak3, and T844M, R653H, V670A mutants were generated Platinum-E retroviral packaging cells^30^ using PEIPro transfection reagent (PolyPlus, New York, NY). 10μg of MSCV plasmid were cotransfected with 4 μg pECO plasmid (Clontech, Mountain View, CA). Plasmid DNA was mixed with polyethyleneimine (PEI) at a 2:1 PEI/DNA ratio and applied to 3 × 10^6^ cells in a 10-cm plate. The medium was changed after overnight incubation, and virus-containing supernatant was collected 48 hours after medium change. Wild-type pro-B cells were infected by spinoculation with 1 mL of viral supernatant containing 10ul of polybrene at the concentration of 10mg/ml. Cells were centrifuged at 3000 × g for 2 hours at 30°C and allowed to recover for 24 hours before medium change and infection was confirmed by flow cytometric analysis for GFP.

### Cell Culture

Wild-type and Jak3 mutant-infected pro-B cell lines used in this study were cultured in IMDM (Wisent, QC, Canada) containing 10% FBS (Wisent), 1X penicillin/streptomycin/L-glutamine (Lonza, Shawinigan, QC, Canada), and 5 x 10^-5^ M β-mercaptoethanol (Sigma-Aldrich St. Louis, MO). Media also contained 5% or 0.5% conditioned medium from the IL-7 producing cell line J558-IL-7^31^. Cell lines were maintained in 5% CO_2_ atmosphere at 37˚C.

### Availability of data

WES data is available from the Sequence Read Archive, accession SRP136503 (https://www.ncbi.nlm.nih.gov/sra/SRP136503). RNA-seq data is available from the Gene Expression Omnibus, accession GSE112506 (https://www.ncbi.nlm.nih.gov/geo/query/acc.cgi?acc=GSE112506).

### Statistical Analysis

All data reported in this study were graphed as mean ± SEM. Statistical analysis was performed using Prism 5.0 (Graphpad Software, La Jolla, CA) using ANOVA or Student’s *t* test. Values with *p* ≤ 0.05 were considered significant, where *p* ≤ 0.05 (*), *p* ≤ 0.01 (**), *p* ≤ 0.001 (***), *p≤* 0.0001 (****).

## Results

### Deletion of genes encoding PU.1 and Spi-B leads to B-cell acute lymphoblastic leukemia

We recently reported that deletion of the genes encoding both PU.1 and Spi-B in B cells under control of the *Cd79a* (*Mb1*) promoter (*Mb1*^+/Cre^ *Spi1*^lox/lox^ *Spib*^-/-^ mice, abbreviated to Mb1-CreΔPB) resulted in a severe impairment to B cell development at the large pre-B to small pre-B cell transition in the bone marrow of 6-10 week-old mice^16^. Mb1-CreΔPB mice that express Cre recombinase and are deleted for Spi-B but are homozygous for the wild type *Spi1* allele were fertile and healthy. In contrast, Mb1-CreΔPB mice had a median survival of 18 weeks, at which point they required euthanasia due to signs of illness, including lethargy and laboured breathing (**Fig. 1A, 1B**). Dissection of euthanized mice revealed enlargement of the spleen and thymus (**Fig. 1A, 1C, 1D**). Histological analysis revealed that normal spleen and thymus organization was completely effaced in moribund Mb1-CreΔPB mice compared to the controls (**Fig. 1E**).

Flow cytometry analysis was used to determine the stage in which leukemic cells begin to infiltrate the spleen and thymus of pre-leukemic Mb1-CreΔPB mice. We observed the presence of B220^+^CD19^+^ cells in the spleen and thymus of Mb1-Cre ΔPB mice aged 11 weeks, but not in either C57BL/6 or Mb1ΔB mice **(Fig. 2A and 2B)**. At the time of euthanasia of leukemic Mb1-Cre ΔPB mice aged >15 weeks there were high frequencies of B220^+^CD19^+^ cells in both organs **(Fig. 2A and 2B)**. Further analysis seeking to determine the phenotype of cells infiltrating the spleen and thymus showed that these organs contained high frequencies of B220^+^ and CD19^+^ cells that also expressed c-Kit, CD43, BP-1, and IL-7Rα (**Fig. 2D** and data not shown). This analysis suggested that these cells were a pre-B cell-like acute lymphoblastic leukemia similar to that previously reported in CD19-CreΔPB mice^17^ or mice deleted for genes encoding PU.1 and IRF4/8^32^. Next, leukemic cells obtained from the spleen or thymus of Mb1-CreΔPB mice were characterized to determine expression of cell surface IgM and Igκ. Analysis of leukemias from Mb1-CreΔPB mice revealed that at least 74% of individual mouse leukemias did not express IgM or Igκ at the cell surface (**Fig. 2E, bottom; 2F, left panel**). 26% of leukemias from Mb1-CreΔPB mouse spleen or thymus had cell surface expression of both IgM and Igκ (**Fig. 2E, top; 2F, left panel**). In contrast, analysis performed on CD19-CreΔPB mice showed that 78% of leukemias had IgM and Igκ on the cell surface **(Fig. 2F, right panel)**. Most leukemias isolated from Mb1-CreΔPB mice had *IgH* rearrangements, although fewer leukemias expressed IgM at the cell surface **(Fig. 2C)**. In summary, these results show that leukemia cells begin to appear in thymus and spleen of Mb1-CreΔPB mice after 11 weeks of age. All leukemias in Mb1-CreΔPB mice at 11-26 weeks of age expressed IL-7R, but most did not express Ig on their cell surface, suggesting that these cells resembled pro-B or large pre-B cells.

**Fig. 1.**
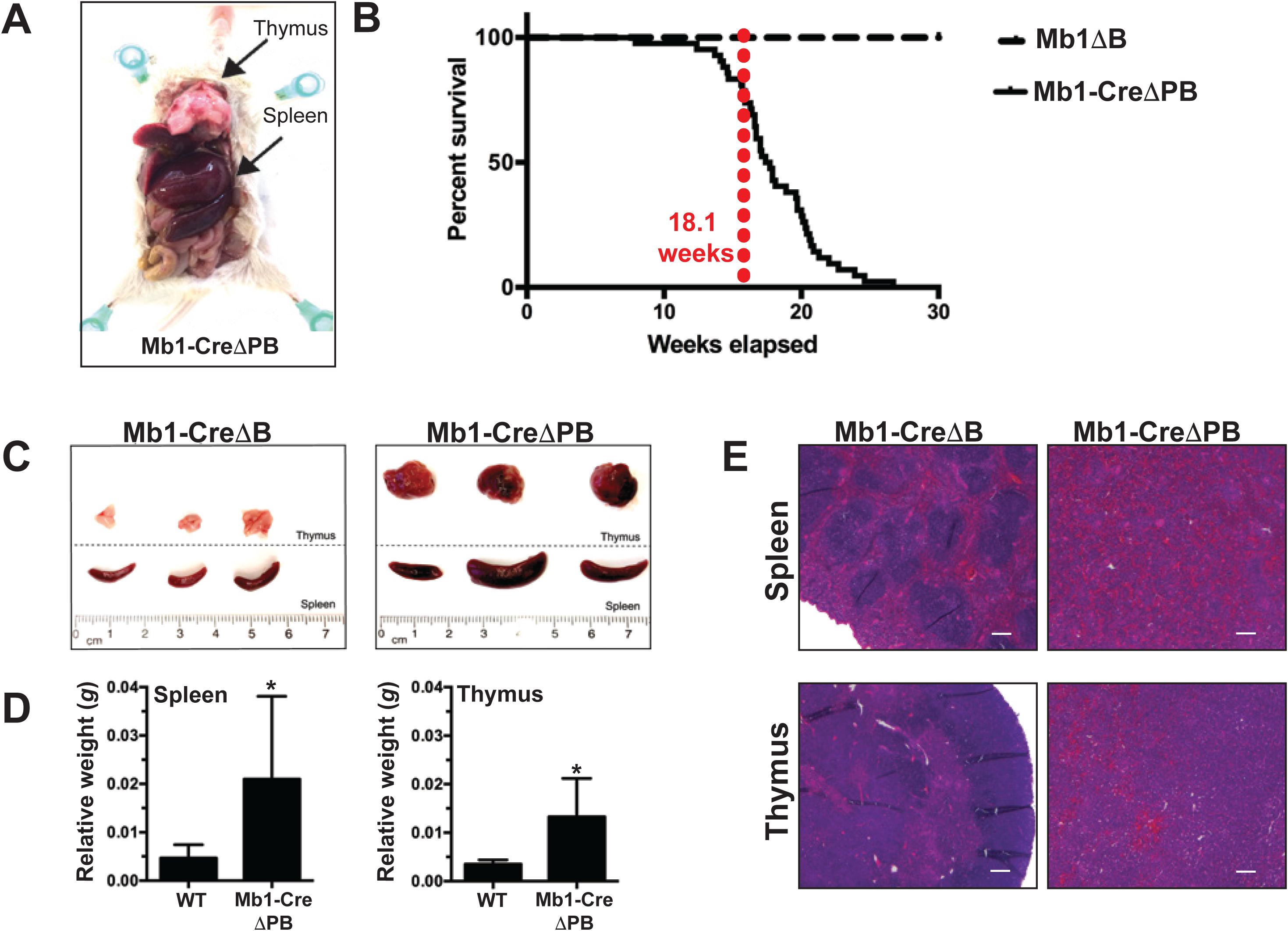
Mb1-CreΔPB mice develop B cell acute lymphoblastic leukemia (B-ALL) **(A)** Mb1-CreΔPB mice (ΔPB) mice developed B-ALL characterized by splenic and thymic enlargement (indicated by arrows). **(B)** Percentage survival of mice of the indicated genotypes: *Mb1^+^*^/Cre^*Spi1*^lox/lox^ *Spib*^-/-^ (Mb1-CreΔPB, n= 43); *Mb1^+^*^/Cre^*Spi1*^+/+^ *Spib*^-/-^ mice (Mb1-CreΔB, n=36) and *Mb1^+^*^/+^*Spi1*^lox/lox^*Spib*^-/-^ (Mb1-CreΔB, n=14). **(C)**. Comparisons of enlarged spleens and thymuses extracted from Mb1-CreΔPB mice compared to control ΔB mice (*Mb1*^+/+^ *Spi1*^lox/lox^ *Spib*^-/-^). **(D)** Spleen (left) and thymus (right) weight in grams (*g*) relative to the body weight in WT and Mb1-CreΔPB mice. WT, n=10 (spleen), 5 (thymus); Mb1-CreΔPB, n=11 (spleen), 8 (thymus). *p≤* 0.05 (*), mean ± SEM. **(E)**. Histologic sections (Hematoxylin & Eosin staining) of spleen and thymus illustrating the lymphocytic infiltration and loss of organs normal structure in Mb1-CreΔPB compared to controls Mb1ΔB (4x).

**Fig. 2.**
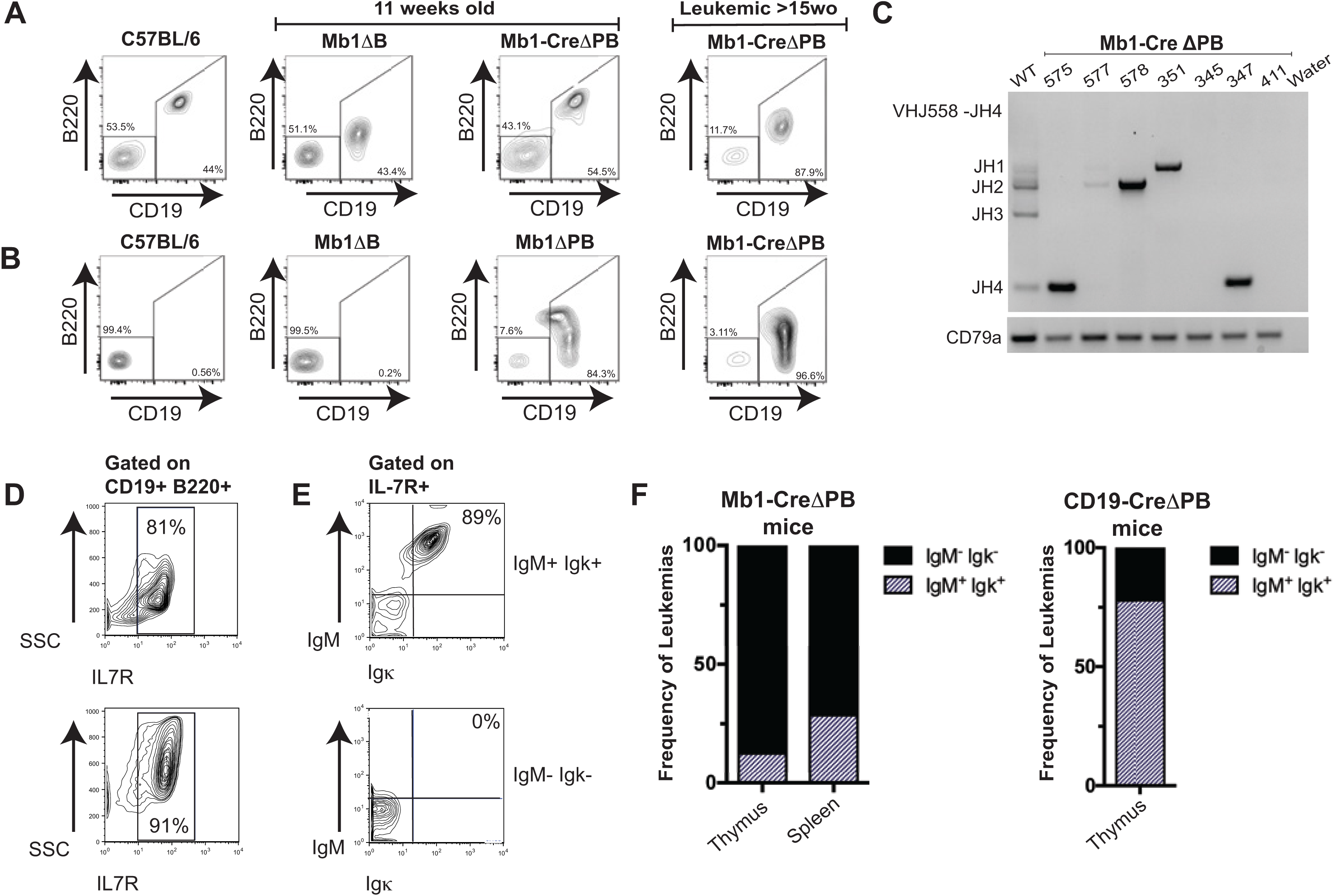
Most leukemias from Mb1-CreΔPB mice resemble pro-B cells and do not express either IgM or Igk at the cell surface. **(A)** Representative flow cytometric analysis for the presence of CD19^+^ B220^+^ B cells in the spleen of C57BL/6 mice; 11 week old Mb1-CreΔB mice and Mb1-CreΔPB mice (*left panels*); and >15 week old Mb1-CreΔPB mice (*right panel*). **(B)** Representative flow cytometric analysis for the presence of CD19^+^ B220^+^ B cells in the thymus of C57BL/6 mice; 11 week old Mb1-CreΔB mice and Mb1-CreΔPB mice (*left panels*); and >15 week old Mb1-CreΔPB mice (*right panel*). **(C)** PCR for detection of heavy chain rearrangements (J558-JH4) in leukemia B cells prepared from the thymus of Mb1-CreΔPB mice (575-411). B cells prepared from WT mouse (C57BL/6) were used as control. The *Cd79* gene was used as control for DNA quality. **(D)** Representative flow cytometric analysis of leukemic cells from Mb1-CreΔPB mice gated on CD19^+^ B220^+^ cells showed that leukemias expressed IL-7R on the cell surface (top and bottom). **(E)** Representative IL-7R^+^ leukemias expressing IgM and Igκ, (top, Ig^+^) and those not expressing IgM and Igκ(bottom, Ig^-^). **(F)** Percentage of leukemias expressing IgM and Igκ(Ig^+^) or not expressing IgM and Igκ(Ig^-^). Left panel, Mb1-CreΔPB mice, n=14 (spleen) and n=8 (thymus). Right panel, CD19-CreΔPB mice, n=9 (thymus).

### Whole-exome sequencing (WES) identifies somatic nucleotide variants in leukemias generated in the lack of Spi-B and PU.1

As described above, Mb1-CreΔPB mice had a variable time to requirement of euthanasia as well as heterogeneity in Ig expression status of leukemias (**Fig. 1 and 2**). This variability suggested that secondary driver mutations are required to induce leukemia in addition to the initiating lesion of *Spi1*/*Spib* deletion. In order to discover potential driver mutations in Mb1-CreΔPB mice leukemias, we performed whole-exome sequencing (WES) analysis of three mouse tumours, 853, 854, and 857, as well as on matched genomic tail DNA as a control **(Supplementary Fig. 1,** data available in **Supplementary Information**). Single Nucleotide Variants (SNVs) for each one of the samples were identified by comparing leukemia and control tail DNA exome sequences using the Strelka somatic variant caller^26^. The number of SNVs identified by Strelka differed for each one of the tumours, where sample 854 showed the highest number of SNVs (3,887) (**Fig. 3A-C**). Samples 853 and 857 presented a very similar number of SNVs, totaling 2,558 and 2,465, respectively. Although the tumours had a variable number of SNVs, the distribution of the SNVs by chromosome followed a similar pattern in which chromosome 2 showed the highest frequency of SNVs, followed by chromosome 11, chromosome 1 and chromosome 7 (**Supplementary Fig. 2A-C**). The majority of the nucleotide alterations in the three tumours analyzed were C•G -> A•T transversions (**Supplementary Fig. 2D**). In order to gain insight into the nature of the mutational processes in Mb1-CreΔPB mice, we determined the mutational signature of leukemias using DeconstructSigs^33^.This package compares the similarity of each tumour sample to a list of previous published signatures generated by the analysis of 40 distinct types of human cancer^34^. This analysis confirmed enrichment of C•G -> A•T transversions in leukemias 853, 854 and 857 (**Supplementary Fig. 3A-C**). Mutational signature analysis performed using DeconstructSigs indicated that the signatures in leukemias 853, 854, and 857 was most similar to COSMIC Signatures 18, 24, 4, and 9 (**Supplementary Fig. 3A-C**). Signature 18, which showed the highest inferred weight for all leukemia samples analyzed, has been also commonly observed in human childhood cancers including B-ALL and neuroblastoma, but does not have a known mechanism^34^. We also observed an enrichment of C->A transversions flanked 5’base by either adenine (A), cytosine (C), guanine (G) or thymine (T) nucleotides. These transversions were consistently flanked at the 3’ base by an adenine (A) nucleotide (**Supplementary Fig. 3D)**.

**Fig. 3.**
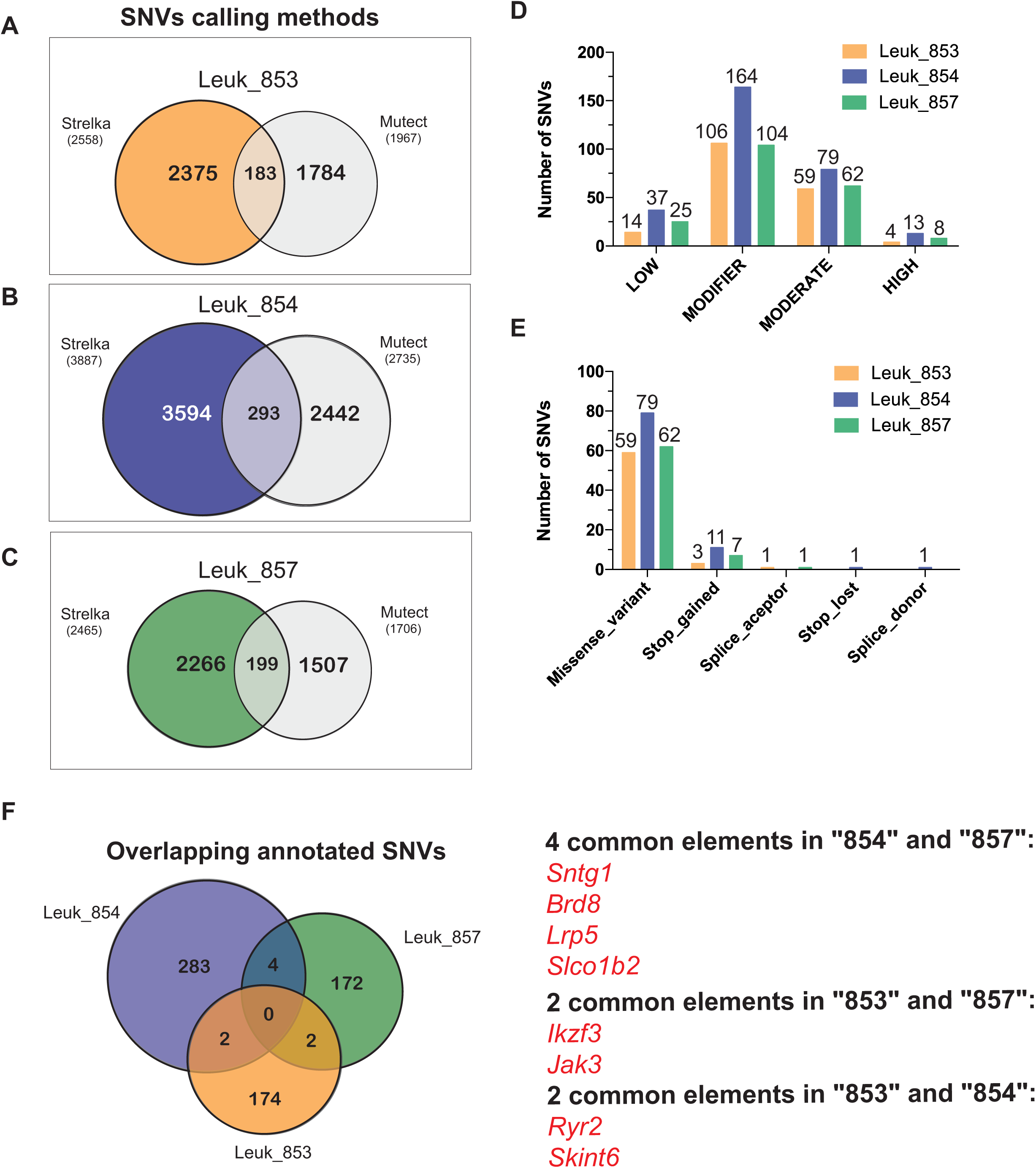
Identification of High Confidence Somatic Nucleotide Variants (SNVs) in Mb1-CreΔPB mouse leukemias. **(A, B and C)** SNVs identified by two different variant caller methods, Strelka and Mutect, were combined and overlapping SNVs were classified as High Confidence SNVs. **(D)** Classification of High Confidence SNVs for impact. Graph shows the number of SNVs classified according to impact. **(E)** Predicted biological effect of the High Confidence SNVs classified as having MODERATE and HIGH impact. **(F)**. Venn diagram showing the overlap among High Confidence SNVs identified in the three individual mouse leukemias.

In order to increase the confidence in the SNVs identified by Strelka and to reduce the number of SNVs without a potential biological function, somatic variants were called with a second variant discovery method, Mutect^27^. Mutect detected fewer SNVs than Strelka (**Fig. 3A-C**). The overlap of SNVs identified by both methods was used to determine high confidence SNVs. We found that by using this strategy the number of SNVs was reduced, resulting in 183 SNVs for sample 853, 293 SNVs for sample 854, and 199 SNVs for sample 857. High confidence SNVs were annotated using the mouse genome (*mm10*) using the SnpEff tool, which also enables the prediction of the biological effect of the variants^28^. We found that the majority of the high confidence SNVs were classified by as having or a MODIFIER or MODERATE deleterious impact while a few SNVs were classified as HIGH (**Fig. 3D**). We next looked at the effect of SNVs classified as MODERATE and HIGH, and found that most of the SNVs classified as MODERATE resulted in missense variants; while the few SNVs predicted as having HIGH impact caused the gain or loss of stop codons, or alterations in splice acceptor or donor sites, defined by the two bases before exon start or ends, respectively (**Fig. 3E)**. We next investigated whether there were common SNVs among the three mouse tumours sequenced. Samples 854 and 857 showed common SNVs in the genes *Sntg1*, *Brd8*, *Lrp5* and *Slco1b2* **(Fig. 3F)**. Samples 853 and 854 showed common SNVs in the genes *Ryr2* and *Skint6*. Finally, samples 853 and 857 showed common SNVs in the genes *Ikzf3* and *Jak3* (**Fig. 3F)**. Interestingly, *Ikzf3* and *Jak3* are well-characterized driver mutations in human pre-B cell acute lymphoblastic leukemia^35^. The identification of common *Ikzf3* and *Jak3* SNVs suggests that this approach is effective at identifying potential driver mutations.

### Mutations in Janus Kinase 1 and 3 (*Jak1*/*Jak3*) and Aiolos (*Ikzf3*) genes are potential secondary drivers of leukemogenesis in Mb1-CreΔPB mice

As cancer driver genes would be expected to contain mutations and be expressed^21^, we also performed RNA-sequencing analysis on leukemias 853, 854, and 857 in order to assess gene expression levels. To filter data based on levels of gene expression, Fragments Per Kilobase of transcript per Million mapped reads (FPKM) was determined from RNA-seq data using Cufflinks. FPKM was plotted against VAF determined using Strelka^26^ for three mouse leukemias 853, 854, and 857. In each of the three samples there were highly expressed genes that also had high VAF (**Fig. 4A-C**). Importantly, sample 853 had high FPKM and VAF for variants in *Jak3* and *Ikzf3* (**Fig 4A**). Sample 854 had high FPKM and VAF for a variant in *Jak1* (**Fig 4B**). Finally, sample 857 had high FPKM and VAF for variants in *Jak3* and *Ikzf3* (**Fig 4C**). The elevated expression levels added to the high VAF of *Jak1*, *Jak3* and *Ikzf3* genes in Mb1-CreΔPB mice leukemias supports the hypothesis that these variants represent secondary driver mutations.

**Fig. 4.**
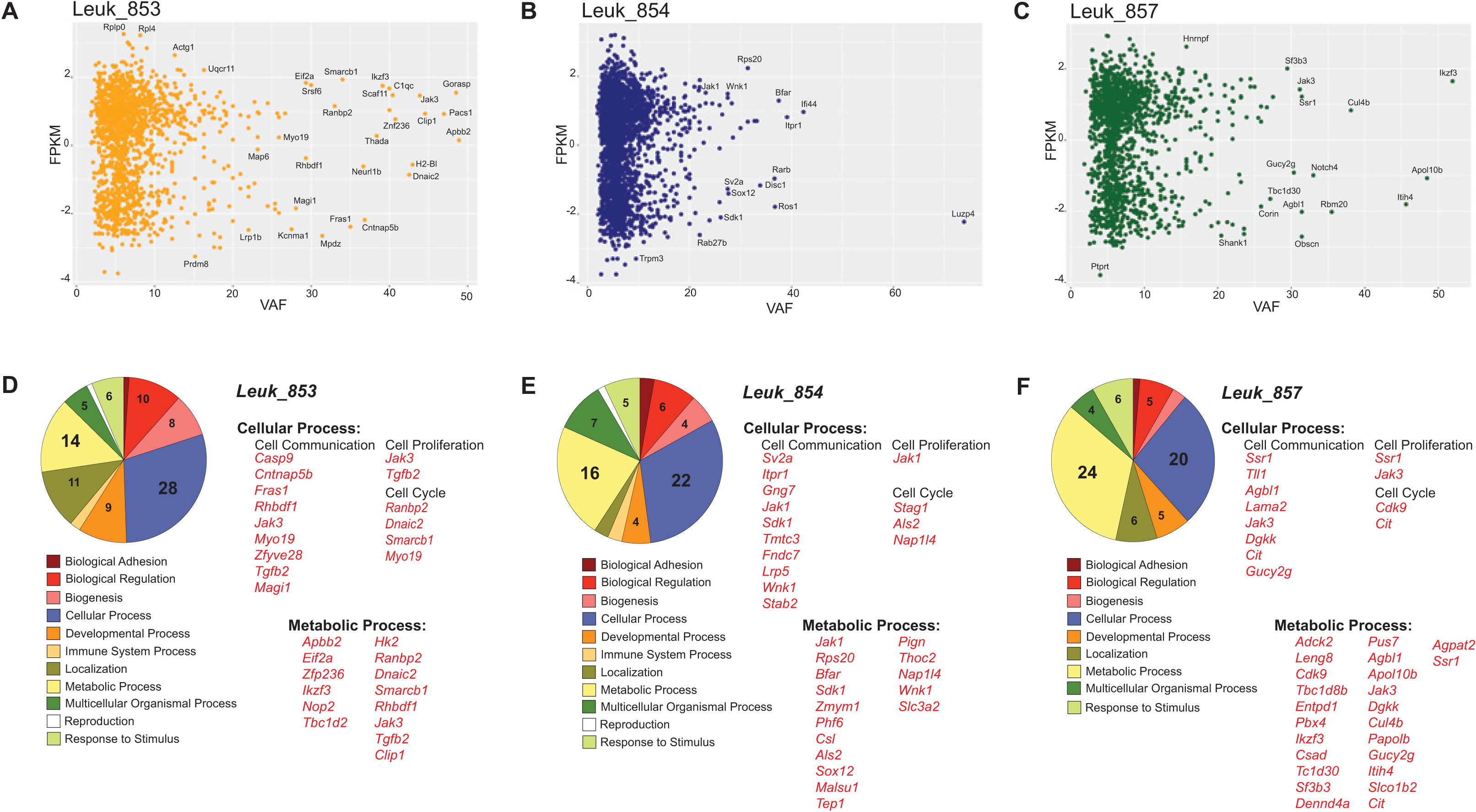
Integration of whole-exome sequencing (WES) and RNA-seq. **(A, B and C)**. Scatter plot correlating the levels of gene expression in Fragments Per Kilobase of Transcript per Million of reads ^mapped (FPKM log10) and the Variant Allele Frequency (VAF) for genes in which FPKM was greater^ than zero. Leukemias 853, 854 and 857 are shown, respectively. **(D, E and F)**. Biological Pathway Analysis in genes with VAF equal or greater than 20% was performed using Panther – Gene List Analysis. Diagram shows the number of genes enriched according the biological process. Enrichment for genes related to “Cellular Process” and “Metabolic Process” is shown for the three samples analyzed.

Next, we sought to define the biological processes in which mutated genes could be involved by performing gene ontology analysis in genes with VAF greater than 20%, using PANTHER Gene List Analysis^36^. We observed a pattern for all three samples where most of the genes were categorized in two major biological processes, “Cellular Process” and “Metabolic Process”. Specifically, genes within “Cellular Process” were subcategorized in sub processes as cell communication, cell proliferation and cell cycle **(Fig. 4D-F)**. Genes as *Cit*, *Cdk9* and *Stag1* are related to the control of cell cycle by regulating cytokinesis, transcription and cohesion of sister chromatids after DNA replication, respectively. Importantly, *Smarcb1* encodes a core subunit protein of the ATP-dependent SWI/SNF chromatin remodeling complex, and was previously identified as a tumor suppressor^37^. *Nap1l4*, encodes a member of the nucleosome assembly protein and has a role as a histone chaperone. Genes involved in cell communication and adhesion, as *Cntnap5b*, *Fras1*, *Sdk1* and *Magi*, were also enriched in our pathway analysis. In summary, *Jak3* and *Ikzf3* mutations were identified as potential drivers of leukemogenesis in Mb1-CreΔPB mice, based on the observation that these genes are mutated in two of three leukemias, have high VAF, and have high levels of expression.

### Recurrent mutations in Janus Kinase 3 (*Jak3*) and Janus Kinase 1 (*Jak1*) in leukemias from Mb1-CreΔPB mice

Analysis of the impact of variants on protein expression showed that SNVs in leukemias 853 and 857 resulted in coding changes in *Jak3* (R653H, V670A, and/or T844M) while the SNV in *Jak1* in sample 854 resulted in a V657F substitution (**Fig. 5A, 5B**). These genes encode for Jak1 and Jak3 that are critically required for signaling through cytokine receptors IL7R and CRLF2^38^. Comparison of Jak3 and Jak1 protein sequences among six different vertebrates showed that amino acids that underwent substitution in consequence of a SNV were highly conserved between these species (**Fig. 5A, 5B**). Sanger sequencing of samples 853, 854 and 857 confirmed the presence of SNVs identified by WES in the *Jak3*, *Jak1* and *Ikzf3*genes (**Supplementary Fig. 3)**. To determine whether these variants represent recurrent leukemia driver mutations, we performed Sanger sequencing on a total of 19 Mb1-CreΔPB leukemias using primers flanking the regions containing the SNVs identified by WES. Interestingly, this examination revealed that Jak3 R653H mutation was detectable in 5/19 leukemias analyzed. *Jak1* mutations V657F or V655L showed even higher recurrence than Jak3 mutation, being detectable in 10/19 leukemias (**Fig. 5D**). *Ikzf3* mutations were detected only in samples 853 and 857, confirming the exome sequencing (**Fig. 5C, 5D**). In summary, these data show that leukemia in Mb1-CreΔPB mice is accompanied by recurrent secondary driver mutations in *Jak1* and *Jak3*.

**Fig. 5.**
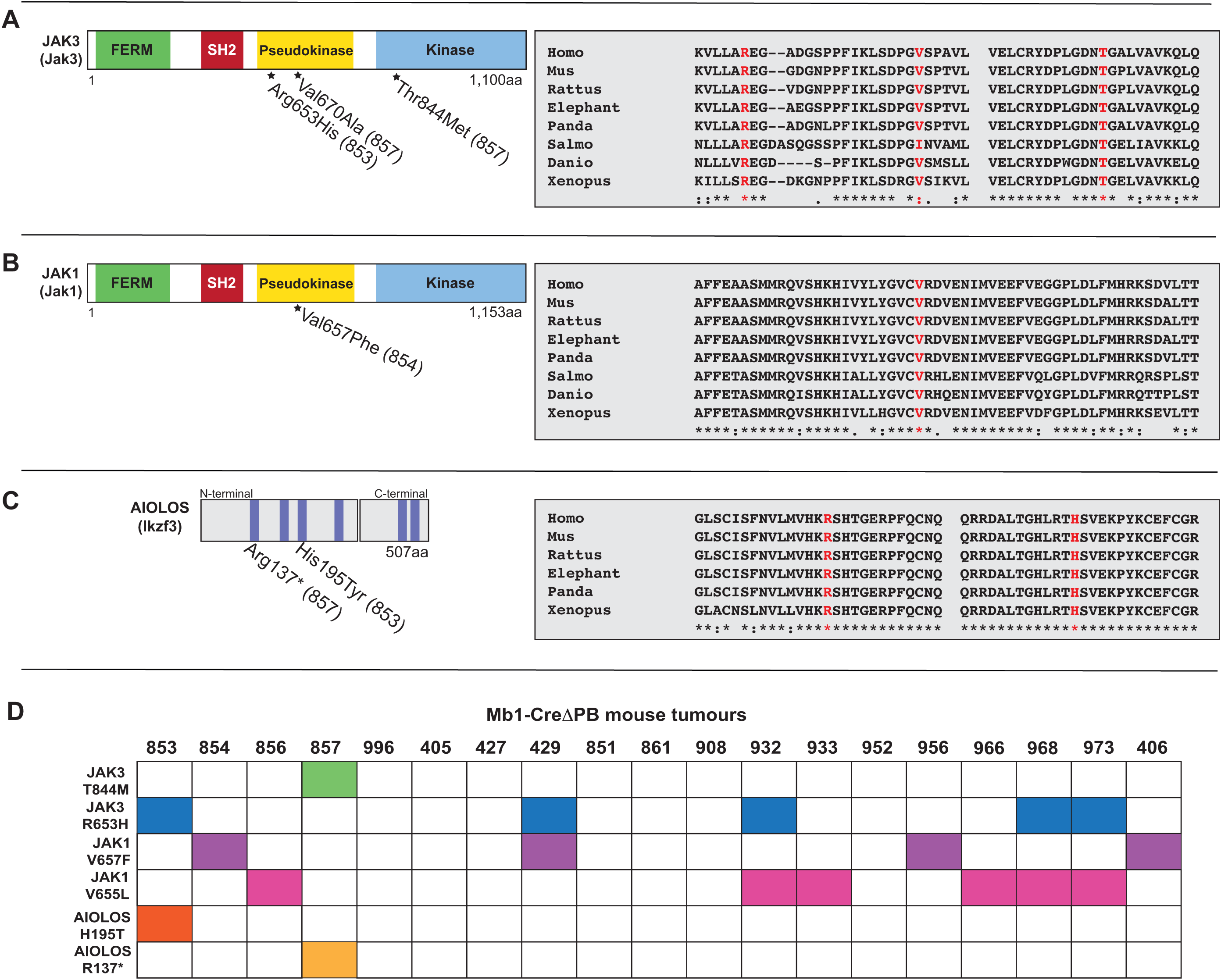
Identified mutations in conserved regions of *Jak3*, *Jak1,* and *Ikzf3* genes. **(A, B and C; *left*)** Schematic shows the protein domains of Jak3, Jak1 and Aiolos (*Ikzf3*). Amino acids substitutions caused by single nucleotide variants identified in the whole-exome sequencing in samples 853, 854 and 857 are also indicated. **(A, B and C; *right*)** BLAST analysis comparing Human (*Homo sapiens*), Mouse (*Mus musculus*), Rat *(Rattus norvergicus),* African Elephant *(Loxodonta africana),* Giant Panda *(Ailuropoda melanoleuca),* Atlantic Salmon *(Salmo salar),* Zebrafish *(Danio rerio),* Tropical Frog (*Xenopus tropicalis*) protein sequences for Jak1, Jak3 and Aiolos shows that amino acids which undergo substitution in consequence of a single nucleotide variation are highly conserved between the two species. **(D)** Summary of Sanger sequencing screening for the presence of amino acids substitutions in Jak1 (V657F and V655L), Jak3 (T844M and R653H), and Aiolos (R137* and H195T) in a panel of nineteen leukemias prepared from Mb1-CreΔPB mice. Filled boxes indicate samples in which mutations were identified by Sanger Sequencing.

### R653H, V670A, and T844M amino acid substitutions in Jak3 confers survival and proliferation advantages

We next tested whether R653H, V670A, and T844M amino acid substitutions in Jak3 were able to confer growth advantage to normal cells. Site-directed mutagenesis was used to introduce these mutations to the wild-type mouse *Jak3* coding region in an MSCV-IRES-GFP vector (a kind gift from Dr. Kevin D. Bunting). Pseudovirus was generated to spin-infect wild-type pro-B cells that grow in cultures containing IL-7 and ST2 stromal cells. Pro-B cells were also infected with MSCV-Jak3 and a MSCV-empty vectors as controls. Following infection, the cell lines were cultured at high (5% conditioned medium) and low IL-7 (0.5%). Flow cytometry was performed every two days for 14 days to determine the frequency of GFP^+^ cells. Pro-B cells infected with *Jak3* mutations R653H, V670A, or T844M outgrew uninfected or control infected pro-B cells at low and high IL-7 concentration **(Fig. 6A, 6B)**. Next, in order to verify whether *Jak3* mutations resulted in greater proliferation, cells infected with mutant Jak3 were plated at the initial concentration of 1x10^5^ cells/ml in conditions of low and high IL-7. Cell numbers and the percentage of viable GFP^+^ cells were determined after cells were kept in culture for the period of four days. At low IL-7 concentration, pro-B cells infected with *Jak3* R653H, V670A, and T844M mutations showed a significant increase in the number of GFP^+^ cells after four days in culture **(Fig. 6C, 6D)**. Therefore, we conclude that T844M, R653H and V670A amino acids substitutions were able to increase the proliferation of pro-B cells in conditions of low IL-7. Taken together, these data indicate that *Jak3* mutations identified in this analysis confer survival and growth advantages to pro-B cells, consistent with a role as driver mutations.

**Fig. 6.**
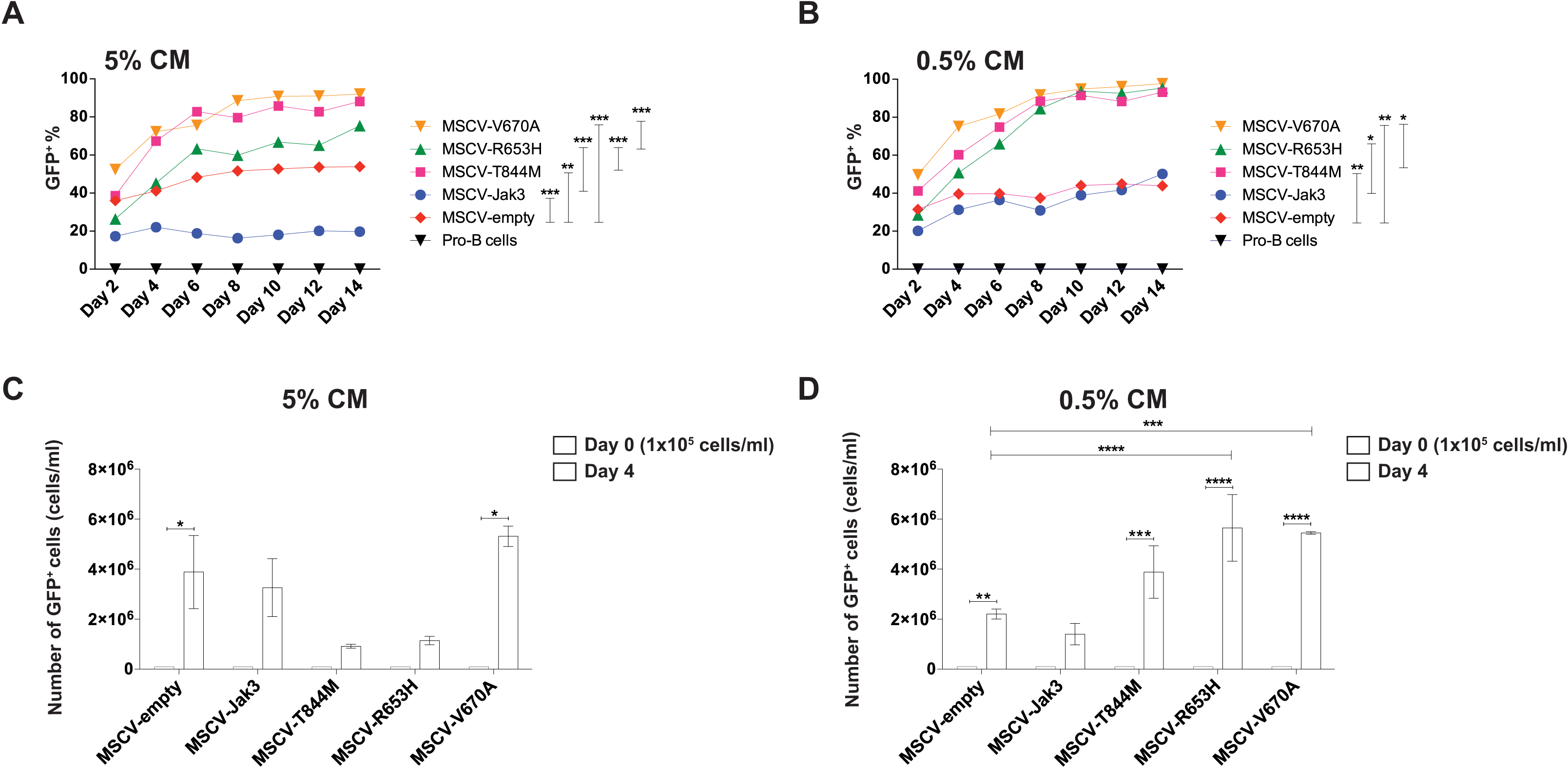
*Jak3* mutations confer growth advantage and increased proliferative potential in normal pro-B cells. **(A and B)** Percentage of GFP^+^ cells over the course of 14 days after infection with MSCV-empty, MSCV-JAK3, MSCV-T844M, MSCV-R653H, MSCV-V670A or MSCV-V670A/T844M in conditions of low (0.5%) and high (5%) IL-7 conditioned medium (CM), respectively. **(C and D)** Absolute number of viable GFP^+^ cells after four days in culture under conditions of low (0.5%) and high (5%) IL-7. Cells were infected with either MSCV-empty, MSCV-JAK3, MSCV-T844M, MSCV-R653H, MSCV-V670A or MSCV-V670A/T844M. Data is presented as number of cells/ml; *n=3*

## Discussion

In this study, we showed that Mb1-CreΔPB mice develop precursor B-ALL that is detectable in mice by 11 weeks of age and results in a requirement for euthanasia in 100% of mice by median 18 weeks. The latency of 11-18 weeks, and the variable frequency of leukemias expressing Ig, show that there is clonal variability, suggesting that secondary driver mutations are required for disease progression. By characterizing the mutation landscape of Mb1-CreΔPB leukemias using whole exome sequencing coupled with RNA-sequencing, we discovered that mutations in *Jak1* and *Jak3* represent recurrent secondary driver mutations. Introduction of mutant *Jak3* into wild type cells was sufficient to promote growth in response to low IL-7 concentrations. Taken together, these results show that deletion of genes encoding PU.1 and Spi-B result in leukemia by predisposition to additional *de novo* driver mutations in genes encoding Janus kinases.

Dysregulation of PU.1 and/or Spi-B expression are known to be involved in human leukemia and have been established as tumor suppressors in mouse models^17,39^. Minimal reductions in PU.1 expression are sufficient to induce a preleukemic condition in hematopoietic stem cells^40^. Whole exome sequencing and whole genome sequencing studies have revealed that inactivating mutations in genes encoding PU.1 and Spi-B are detectable and infrequent in human leukemia^41,42^. In contrast, PU.1 levels are repressed by frequently occurring chromosomal abnormalities such as FLT3 internal tandem duplication (FLT3-ITD)^43^ and RUNX1-ETO fusion^44^. Spi-B levels are repressed by ETV6-RUNX1 chromosomal fusion^39^. It is not known how reduced PU.1 and/or Spi-B might lead to acquisition of secondary driver mutations. However, PU.1 and Spi-B are involved in attenuating IL-7R signaling in developing pro- and pre-B cells. Target genes of PU.1 and Spi-B involved in regulating IL-7-induced proliferation include *Btk* encoding the tumour suppressor Bruton tyrosine kinase^15^, and *Blnk* encoding B Cell Linker Protein^45^. Btk and Blnk work together to attenuate IL-7R signaling in developing B cells^46^. We speculate that mutagenesis in Mb1-Cre-ΔPB is associated with genomic instability due to uncontrolled cell proliferation. Interestingly, a similar disease is induced by combined deletion of PU.1 and IRF4, suggesting a common axis of gene regulation by PU.1/Spi-B and PU.1/IRF4^32^.

We identified a number of single nucleotide variants (SNVs) in our analysis, which suggest that several rounds of mutational events contributed to leukemogenesis. To select the potential leukemia initiator lesions, we focused our attention in genes that were highly expressed and had high variant allelic frequencies (VAF), reasoning that genes within this category are likely to have originated earlier in leukemia progression. This analysis identified Aiolos (*Ikzf3*), Jak1 (*Jak1*) and Jak3 (*Jak3*) as potential secondary driver mutations. These genes are involved in signaling downstream of cytokine receptors during B cell development. Cytokine receptor genes are commonly mutated in human B-cell acute lymphoblastic leukemias^47^, and mutations in *Jak3* and *Ikzf3* have recently emerged as novel mutated genes in high-risk B-ALL^48^. In particular, our analysis focused on R653H, V670A, and T844M mutations in *Jak3*. Jak3 R653H and V670A mutations were recently observed in B-ALL generated in activation-induced cytidine deaminase (AID) deficient mice^49^. Human equivalents of Jak3 R653H and V670A have been described in human ALL (R657Q, V674F), and furthermore have been shown to function by activation of proliferation in response to interleukin-7^50^. Importantly, human JAK3 V674F is sufficient to induce T-cell acute lymphoblastic leukemia when retrovirally delivered to hematopoietic precursors^50^. *Jak1* missense mutations V655L or V657F were detectable in 10 of 19 leukemias in our analysis. Jak1 V657F also corresponds to a human *JAK1* mutation that is a frequent driver in human B-ALL (V658F)^51^. Interestingly *JAK1* V658F mutation has been previously observed in T-cell acute lymphoblastic and acute myeloid leukemias^52^. V658F is also thought to be paralogous to the JAK2 V617F mutation that functions as a driver in more than 95% of cases of polycythemia vera^53^. In summary, the mutations identified in this study in *Jak1* and *Jak3* are relevant to human leukemia.

Taken together, this study confirms that leukemia in Mb1-CreΔPB mice is accompanied by *de novo* recurrent secondary driver mutations in the Janus Kinase signaling pathway. Mutations in the JAK signaling pathway are recurrent in human leukemia including the recently discovered Ph-like classification that represents a high priority for discovering new therapies^9,10^. In order to study the genetic clonal evolution that underlies diseases such as Ph-like leukemia, there is still a need for mouse models. In order to advance understanding, mouse models should have the following characteristics: 1) They should develop leukemias with high penetrance and reproducibility, 2) they should replicate the genetic and molecular heterogeneity of tumors and involve de novo mutations 3) they should occur in immune competent mice, and 4) they should mimic that clinical behaviour of human disease^54,55^. The Mb1-CreΔPB mouse model develops B-ALL with 100% penetrance by 18 weeks of age that is driven by heterogeneous *de novo* driver mutations. This mouse model may be useful to determine the effects of molecular targeted therapies on intratumoral heterogeneity and clonal evolution in B-ALL.

## Acknowledgements

We thank Michelle Ho for assistance with genotyping of mice. We thank Michael Reth (Freiburg, Germany) for providing Mb1-Cre mice. We thank Dr. Kevin D. Bunting (Winship Cancer Institute, Atlanta, USA) for providing the MSCV-IRES-GFP-JAK3 vector. We also thank scientists and staff of the McGill University and Genome Quebec Innovation Centre for performing library construction and next-generation sequencing. We also thank Kristin Chadwick and the London Regional Flow Cytometry Core Facility for assistance with flow cytometric analysis.

---

**Supplementary Fig. 1.**
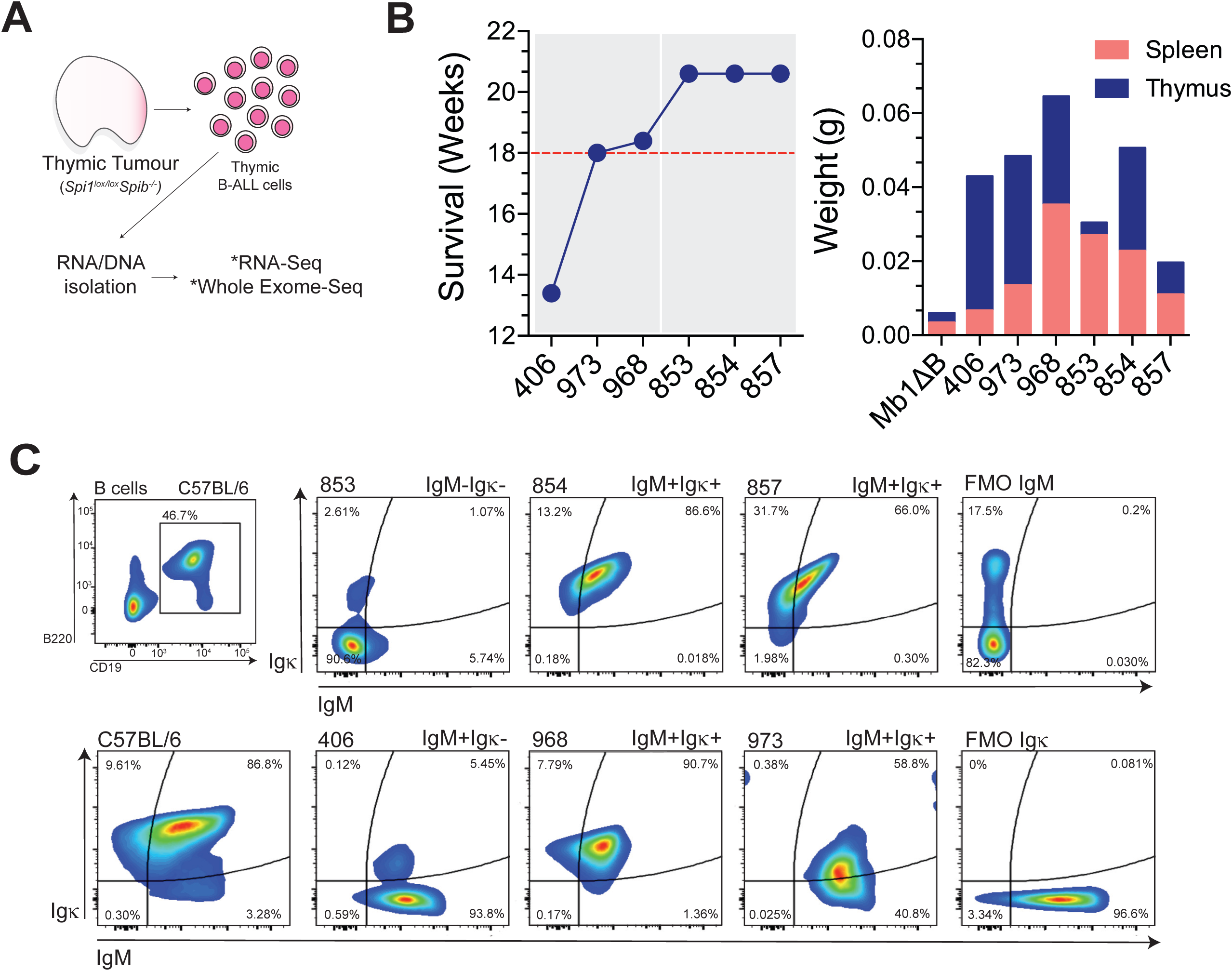
Experimental design for RNA-seq and WES-seq experiments. **(A)** DNA and RNA were isolated from leukemic cells extracted from thymus of leukemic mice. (**B)** Weeks of survival and weight of the organs extracted from representative leukemic Mb1-CreΔPB mice. Samples 853, 854, and 857 were used for sequencing experiments. (**C)** Flow cytometric characterization (IgM, x-axis; Igκ, y-axis; gated on CD19^+^ B220^+^ cells) of leukemic mice in which DNA and RNA were isolated for sequencing. C57BL/6, control; 853, 854, 857, leukemic Mb1-CreΔPB; FMO = fluorescence minus one.

**Supplementary Fig. 2.**
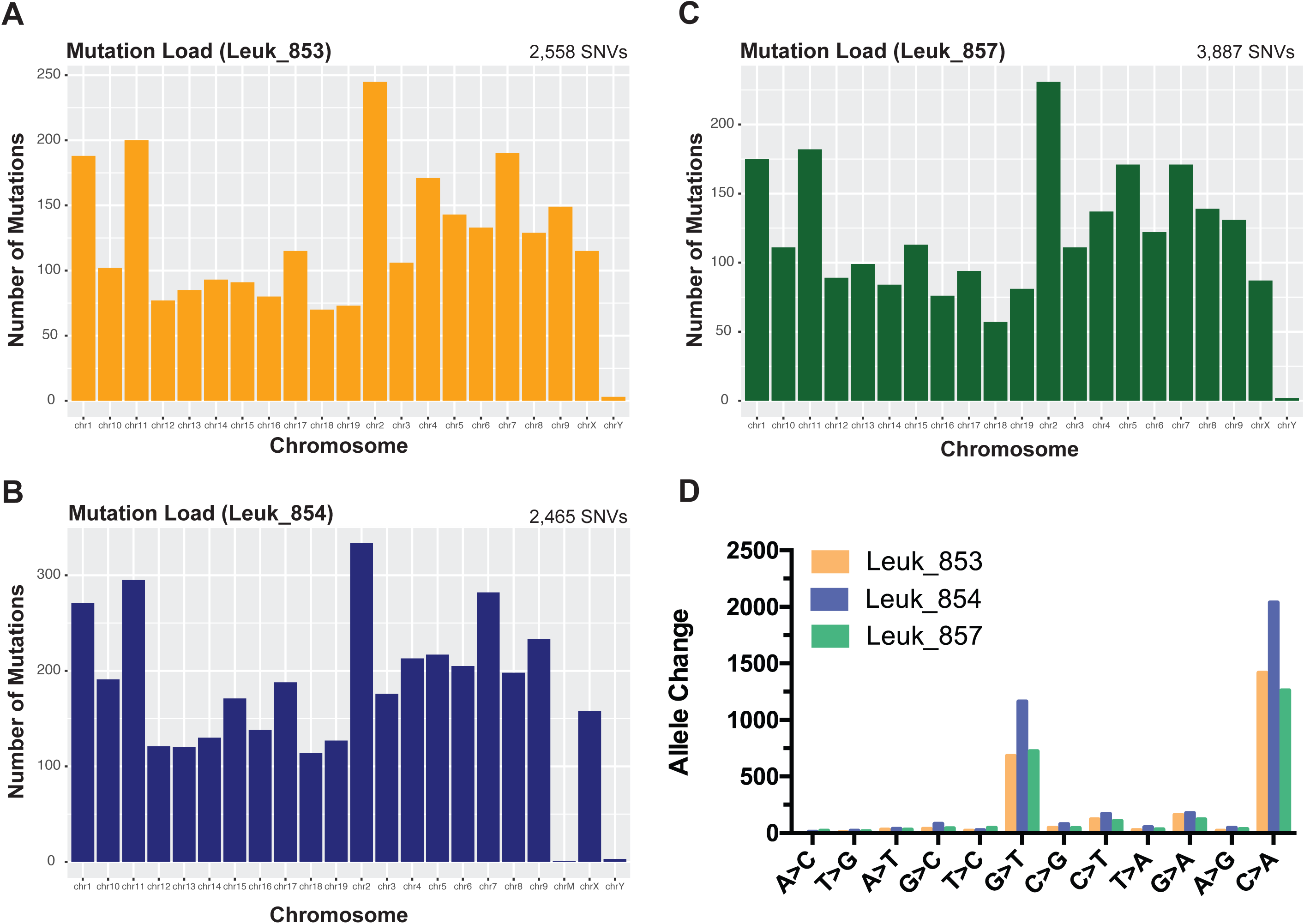
Characterization of the nucleotide somatic nucleotide variants (SNVs) identified by Strelka somatic variant caller in three Mb1-CreΔPB mice leukemias. **(A, B and C)** SNV distribution according to chromosome number in leukemias 853, 854 and 857, respectively. **(D)** Allelic change frequency for 853, 854 and 857 leukemias showing an enrichment for G>T and C>A substitutions.

**Supplementary Fig. 3.**
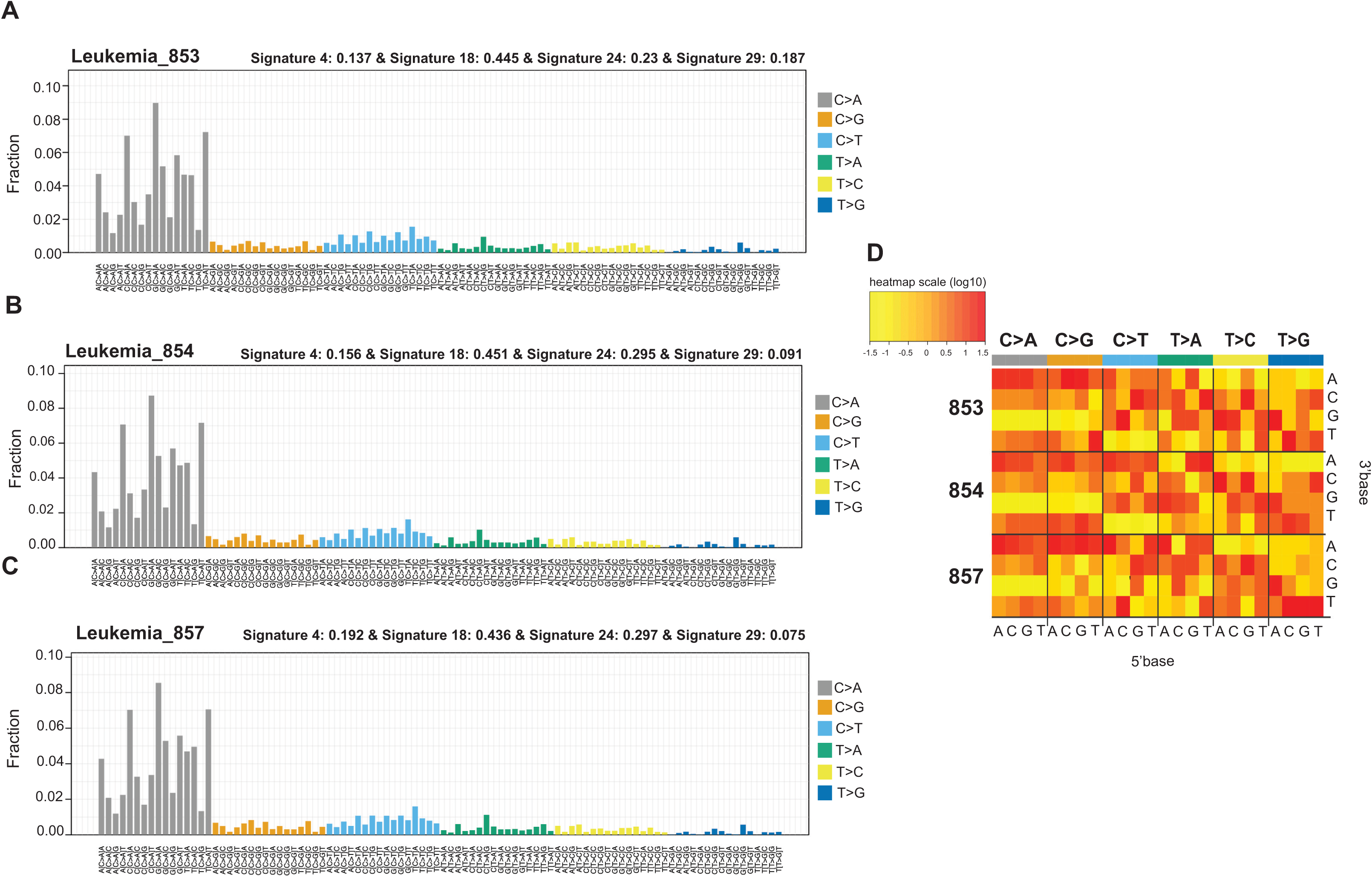
Mutational signature of Mb1-CreΔPB mouse leukemias shows strong bias to C•G -> A•T transversions. **(A-C)** DeconstructSigs reconstruction analysis of mouse leukemias 853, 854 and 857. COSMIC mutation signatures were used to reconstruct the mutational profile of mouse leukemias and to predict similarity. **(D)** Heat map illustrating the mutational context of 5’ and 3’ nucleotides surrounding the transitions identified in samples 853, 854 and 857.

**Supplementary Fig. 4.**
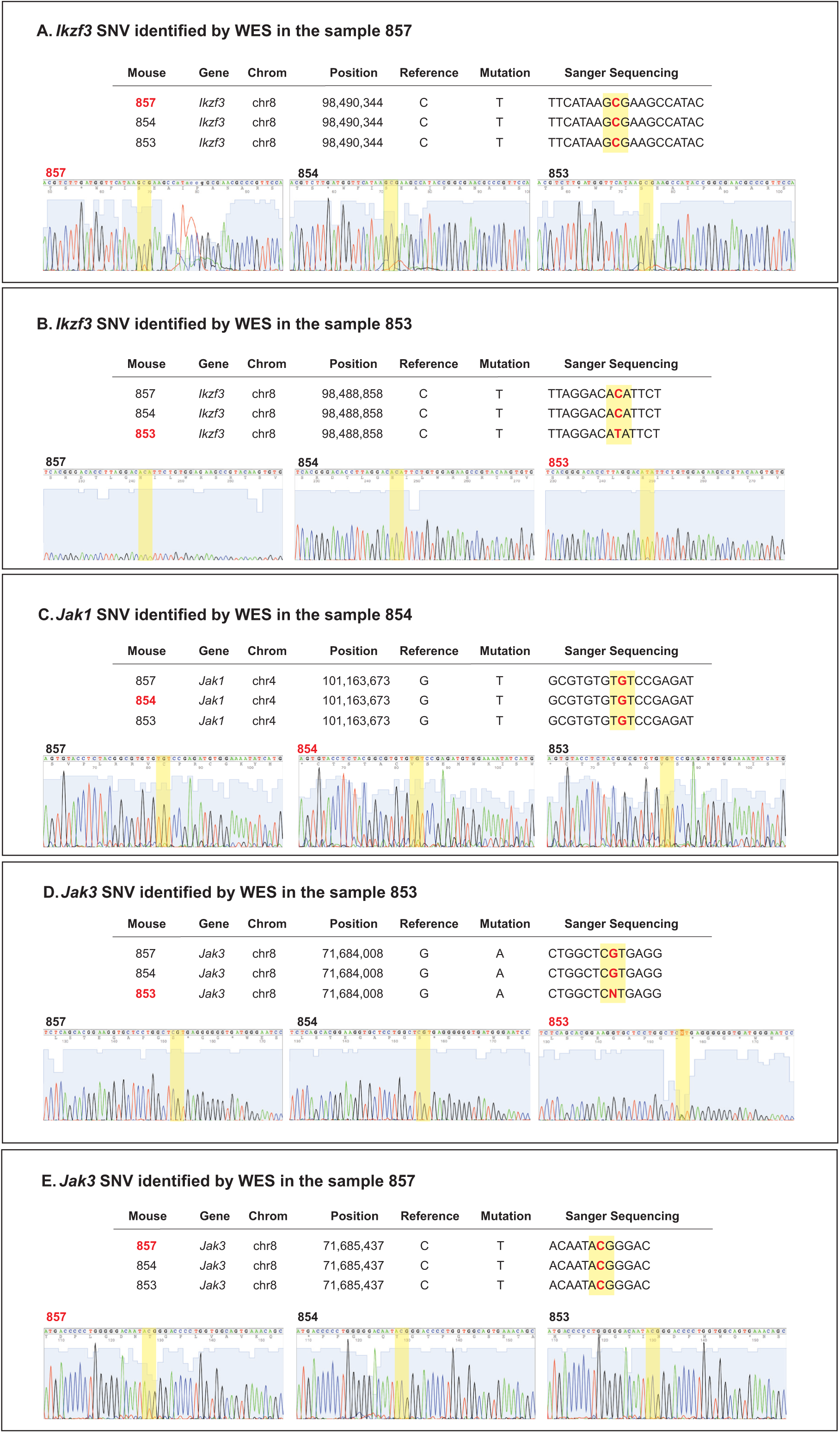
Sanger sequencing confirms the presence of SNVs identified by WES. PCR for amplification of the region containing the SNVs were performed on cDNA synthesized from RNA prepared from mouse tumours. PCR products were submitted for Sanger sequencing using primers targeting the region of interest. **(A)**; Sanger sequencing results of region containing *Ikzf3* SNV identified in the sample 857, transition C-T; **(B)** *Ikzf3* SNV identified in the sample 853, transition C-T; **(C)** *Jak1* SNV identified in the sample 854; transition G-T; (**D**) *Jak3* SNV identified in the sample 853; transition G-A. (**E**) *Jak3* SNV identified in the sample 857; transition C-T. Chromatograms are shown for each SNV investigated. Note that most SNVs are detectable at 50% or lower allelic frequency.

